# Solid State High Throughput Screening Microscopy

**DOI:** 10.1101/602011

**Authors:** M. Ashraf, S. Mohanan, B Sim, A. Tam, D. Brousseau, S. Thibault, A. Corbett, G. Bub

## Abstract

We introduce a solid state high throughput screening (ssHTS) imaging modality that uses a novel Newtonian telescope design to image multiple spatially separated samples without moving parts or robotics. Conventional high-throughput imaging modalities either require movement of the sample to the focal plane of the imaging system^1–3^ or movement of the imaging system itself^4,5^, or use a wide-field approach to capture several samples in one frame. Schemes which move the sample or the imaging system can be mechanically complex and are inherently slow, while wide-field imaging systems have poor light collection efficiency and resolution compared to systems that image a single sample at a given time point. Our proposed ssHTS system uses a large parabolic reflector and an imaging lenses positioned at their focal distances above each sample. A fast LED array sequentially illuminate samples to generate images that are captured with a single camera placed at the focal point of the reflector. This optical configuration allows each sample to completely fill a sensors field of view. Since each LED illuminates a single sample and LED switch times are very fast, images from spatially separated samples can be captured at rates limited only by the camera’s frame rate. The system is demonstrated by imaging cardiac monolayer and *Caenorhabditis elegans* (*C. elegans*) preparations.

---

Our current prototype (Fig 1) uses a 500 frames/second machine vision CMOS camera and a commercially available 256 element LED array controlled by an Arduino microcontroller, which can switch between LEDS at 16 KHz. A single-element 100 mm focal length, 25mm diameter plano-convex lens is placed above each sample so that collimated light is projected to a 100mm focal length parabolic reflector, which then creates an image on the detector. The high light collection efficiency inherent in this design allows images to be captured with sub 200 microsecond exposure times. The camera is synchronised with the LED array via a TTL signal from the microcontroller so that a single frame is captured when any LED is on. This setup can rapidly switch to image any dish under the parabolic reflector without moving the sample or camera. In addition, the system can acquire data from several dishes near-simultaneously by trading-off the number of samples for frame rate: for example, 50 dishes can be captured at 10 fps, or any two dishes can be recorded at 250 fps.

**Figure 1.**
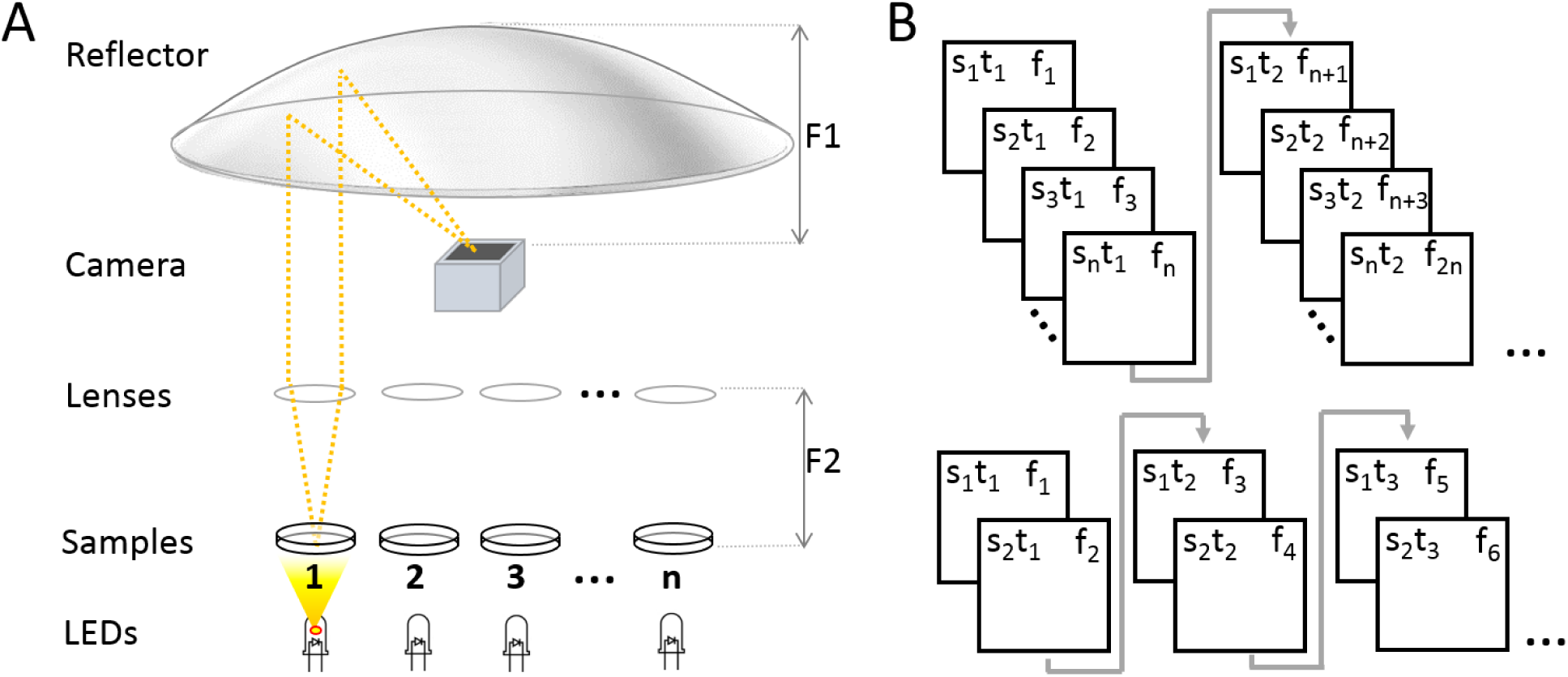
A) The solid state high throughput imaging system uses a parabolic reflector to image samples directly on a fast machine vision camera (acA640-750um - Basler ace), located at the focal point of the reflector (F1). Single element plano-convex lenses are used as objectives, with samples positioned at their focal point (F2). Samples are sequentially illuminated using a fast LED array (Adafruit Industries Dotstar RGB LED 242mm diameter disk) controlled by an Arduino microcontroller: a sample is only projected on the sensor when its corresponding LED is ‘on’. B) (top) Sample *s*, is captured at time *t*, on frame *f*. For a total of *n* samples, each sample is captured once every *n* frames; (bottom) a smaller subset of samples can be imaged at higher temporal resolution by reducing the number of LEDs activated by the microcontroller through software.

The high numerical aperture and large field of view offered by parabolic mirrors have made them very attractive to imaging applications beyond the field of astronomy. However, parabolic mirrors introduce significant off-axis aberrations which corrupt any widefield image formed^6,7^. This has resulted in compromises, such as restricting imaging to the focal region and then stage-scanning the sample^8^ which have limited its use to niche applications. In our design, transillumination from LEDs far from the sample and collimation from the imaging lens results in mostly collimated light being refocused by the parabolic mirror, thereby avoiding the introduction of significant aberrations.

Propagation-based phase contrast in our imaging system is generated when collimated light from the LED passes through and is diffracted by the sample. Light which remains in the collection cone of the imaging lens is then refocused on the sensor by the parabolic reflector at an oblique angle (Fig 2a). As a result of this angle, the image moves through focus from one side of the detector plane to the other. The region over which the image is in focus is determined by the depth of focus of the parabolic mirror (approximately 5 mm). The distance along the chief ray between the image formed at one side of the detector and the other is given by *D*_*f*_ = *D*_*s*_sin(*θ*), where *D*_*s*_ is the width of the sensor and *θ* is the angle of the chief ray. For our system, *D*_*s*_ is 2.4 mm, and *θ* is always less than 60 degrees, so *D*_*f*_ is always less than 2 mm and the entire image therefore remains inside the Rayleigh length of the parabolic focus.

**Figure 2.**
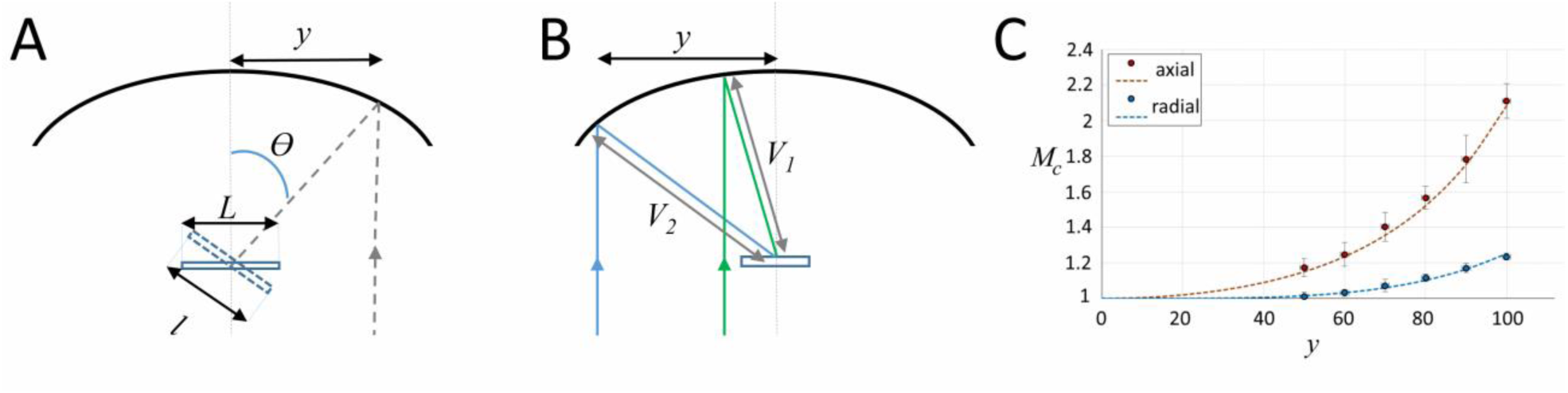
A) The chief ray (dashed line) arrives at the detector plane at an incidence angle *θ* which increases with lateral displacement, *y*. The image is stretched in the direction parallel to *y* by a factor of *L/l*. B) The image is isotopically magnified as the distance between the mirror and the image increases (*V2*>*V1*) as *y* increases. C) The combined magnification, *MC*, shows the impact of the combined transformation on the magnification in both image dimensions (*y*′ parallel to *y*, and *x*′ orthogonal to *y*). Red dots (measured) and dashes (predicted) show magnification in *y*′, and blue dots (measured) and dashes (predicted) show magnification in *x*′. Sample images are shown in the supplement.

Images are subject to two well-defined transformations: (i) a stretch due to the image meeting the camera plane obliquely and (ii) a small variation in magnification as a function of the separation between the optical axes of the objective lens and parabola. These image transformations can be compensated by post-processing the captured images using equations derived from geometric optics as described below.

Light from the sample arrives at the detector plane at an incidence angle, theta, which increases with lateral displacement between objective and mirror axes, *y* (Fig 2a). As the image itself is formed normal to the chief ray, the detector plane captures a geometric projection of the image which is stretched in the direction of *y*. The magnitude of the stretch is given by

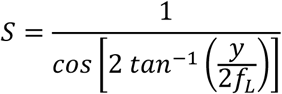

where *S* is the magnitude of the stretch in one axis, *y* is the lateral displacement, and *f*_*L*_ is the focal length of the parabolic mirror. In addition to geometric projection, there is also a small variation in magnification which is the same in both image dimensions (*y*′ parallel to displacement *y*, and *x*′ orthogonal to *y*). This is due to the distance between the parabolic mirror surface (the last focusing element) and the focal point (*V*) increasing as a function of *y* (Fig 2b). The magnification is then given by the ratio of *V* to the focal length of the objective lens, *f*_*L*_. As *V*(*y*) can be calculated precisely for a parabola, it allows us to calculate the magnification, *M*, as a function of *y, f*_*L*_ and mirror focal length, *f*_M_:

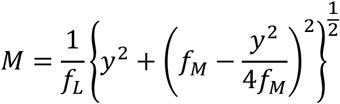

The combined magnification (*M*_*C*_) from global scaling and geometric projection along the *x*′ and *y*′ dimensions is shown together with experimental data in Fig 2c.

We demonstrate the system using two popular biological models that can be positively impacted by new high-throughput screening modalities. Cultured cardiac monolayer preparations, generated either from neonatal animal hearts^9^ or from human pluripotent stem cells^10^, are used to study arrhythmogenesis in controlled settings and are subject to intense research due to their potential for screening compounds for personalized medicine. *C. elegans* are used as model organisms to study the genetics of aging and biological clocks^11^, and, due to highly conserved neurological pathways between mammals and invertebrates, are now used for neuroprotective compound screening^12^. High throughput screening technologies for cardiac culture and *C. elegans* systems are therefore of particular interest to the research community. Figure 3a shows simultaneous recordings from four dishes containing monolayer cultures of neonatal cardiac cells at 40 fps per dish. Here, motion is tracked by measuring the absolute value of intensity changes for each pixel over a six frame window^13^. Intensity vs time plots (Fig 3b) highlight different temporal dynamics in each preparation, and an activation map from one of the dishes shows conduction velocity and wave direction data (Fig 3c). *C. elegans*, plated in 35 mm petri dishes in standard agar medium, can similarly be imaged, here at 15 fps for 4 dishes over a period of 5 minutes (Fig 3d-f). *C. elegans* motion paths (Fig 3d), which are often used to quantify worm behavior, can be extracted from each image series using open-source software packages.

**Figure 2.**
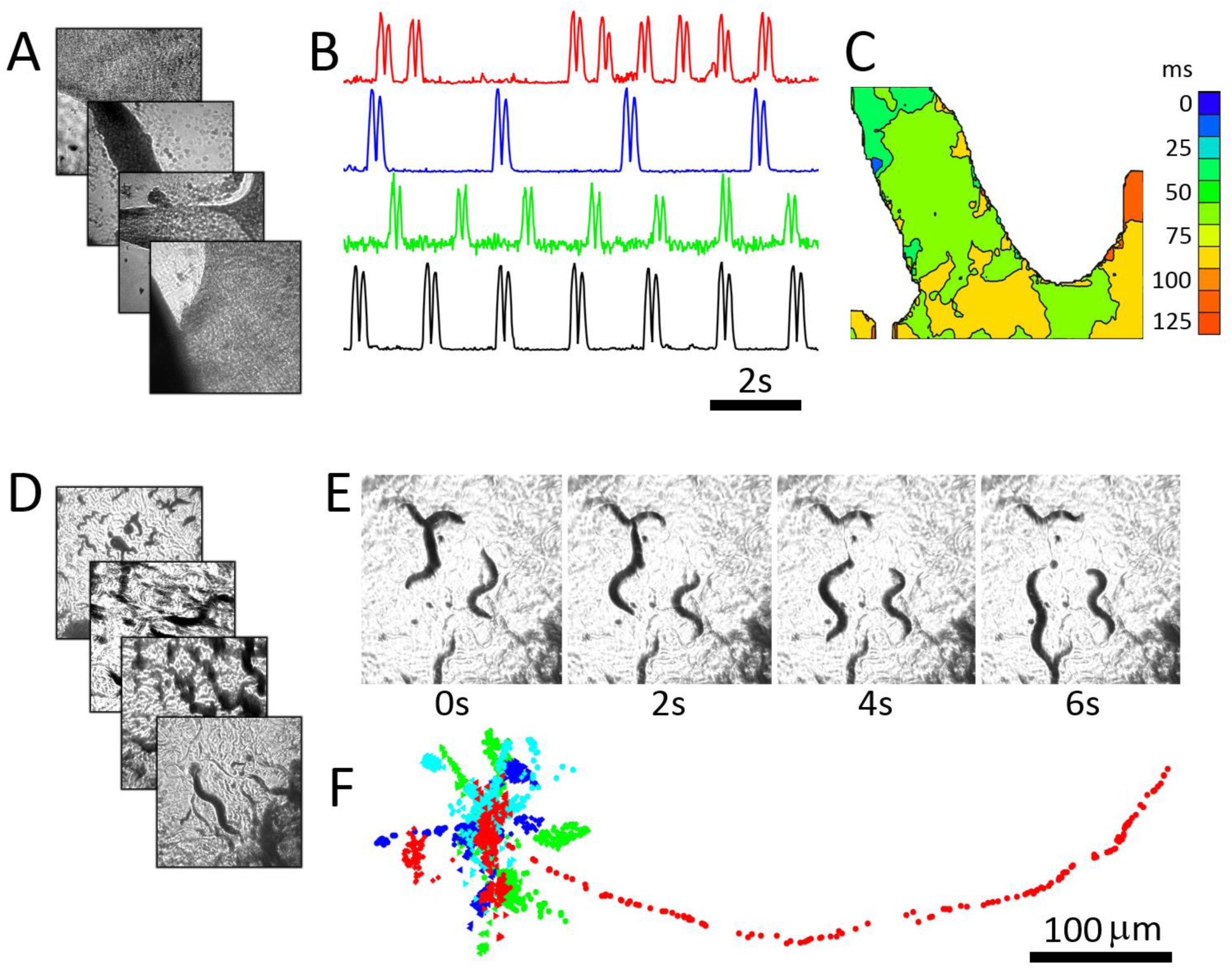
A) Four cardiac monolayer preparations, imaged in four separate petri dishes are imaged at 40 fps/dish. B) Intensity vs time plots obtained from the four dishes show different temporal dynamics; C) an activation map from the second dish (blue trace in A) can be used to determine wave velocity and speed; D) four *C. elegans* dishes imaged simultaneously at 15 fps/dish; E) Images from one dish every 30 frames (2 second intervals) shows *C. elegans* motion; F) The location of five worms in each dish were tracked simultaneously from data recorded at 15 Hz over 250 frames using open source WormTracker software. Dots in different colours (blue, cyan, green, and red) shows the tracked positions from plates 1-4 respectively. Each image in A, D and E shows a 2mm × 2mm field of view.

The push to develop new high throughput screening modalities^14,15^ has resulted in several innovative approaches, ranging from the use of flatbed scanners for slowly varying preparations^16^ to ‘on-chip’ imaging systems which plate samples directly on a sensor^17–21^, to super-resolution methods which allow for high-resolution wide field-imaging by capturing multiple pictures per time point^22^. Despite these advances, methods that accommodate a biologists’ typical workflow - e.g. comparing multiple experimental samples plated in different petri-dishes, have largely depended on automation of conventional microscopes. The performance of these conventional high-throughput screening approaches can be quantified by considering the utilisation of the data rate set by the camera, i.e. camera resolution times frame rate. The central challenge faced by these systems is to maximize the number of pixels that capture data while minimizing the number of pixels lost due to the space between samples in a widefield image or frames lost due to the transit time between individual samples. ssHTS overcomes these constraints by reducing transit time between samples to less than a millisecond without the use of automation or relying on a widefield imaging approach, while allowing for an optimized field of view. In addition, as the capacity of ssHTS is limited only by the size of the parabolic mirror, the system can in principle be scaled up to handle hundreds of samples in a flexible manner.

## Supporting information

Zemax simulations and rescaling results

## Acknowledgements

We would like to thank R.S. Branicky and S. Hekimi (McGill University) for the C.- elegans preparation, A. Caldwell (McGill University) for preparation of cardiac samples, C. Sprigings (McGill University) for programming assistance.

