## Supplementary material for "Solid State High Throughput Screening Microscopy": Zemax simulations and rescaling results

### ZEMAX model of the image formation process

To support the experimental results, a ZEMAX simulation of the experimental design was produced (Figure S1). This simulation used an f=100 mm doublet lens (Thorlabs AC254-100-A) located one focal length from the sample. The sample was configured to be an opaque grid of the same 200 µm pitch and 4 mm × 4 mm area as that used in the experiment. The lateral distance between the optical axes of the lens and the mirror was changed (Table 1). All other parameters remain the same.

To simulate the limited ray angles incident upon the grid from the LED, a 5 mm diameter aperture was placed on the lens. This limited the ray angles to <4 °, which is what would be expected from the LED when situated >50 mm from the 4 mm samples. In both simulated and experimental systems, it is the restriction of the ray angles incident upon the mirror (Edmund Optics #68-793, f=100 mm, f#=0.48) which is essential to achieve high imaging quality of the sample and limit the introduction of astigmatism and other low order aberrations to the image.

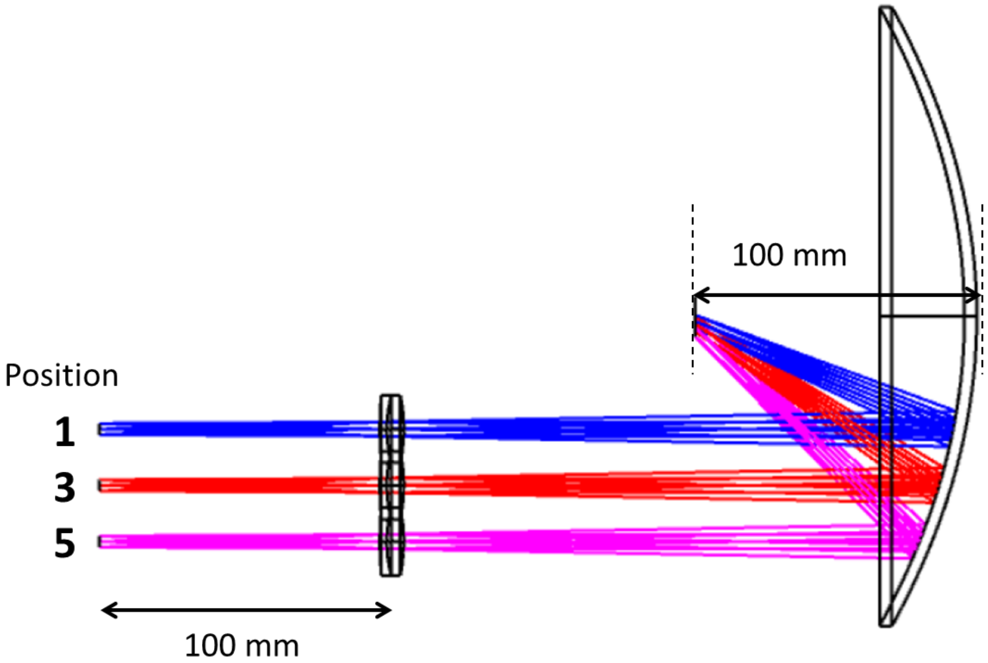


Figure S1: ZEMAX simulation of the ssHTS system. The colour coded rays correspond to the positions of the sample with respect to the optical axis of the mirror. The positions 1-9 correspond to specimens located 10 mm to 90 mm distant from the optical axis in 100 mm steps (Table 1).

Table 1: Location of off-axis lens positions and corresponding angles of incidence of chief ray (see Figure S4).

| Lens position | Distance from mirror axis (mm) | Chief ray angle of incidence (°) |
|---|---|---|
| 0 | 0 | 0 |
| 1 | 10 | 5.7 |
| 2 | 20 | 11.4 |
| 3 | 30 | 17.1 |
| 4 | 40 | 22.6 |
| 5 | 50 | 28.1 |
| 6 | 60 | 33.4 |
| 7 | 70 | 38.6 |
| 8 | 80 | 43.6 |
| 9 | 90 | 48.4 |
| 10 | 100 | 53.1 |

Simulations of the grid images are shown in Figure S2 for off-axis locations of 40-90 mm. It can be seen that there the principal distortion is a vertical scaling. There is also a uniform scaling of the image. Both are present in the experimental data (shown for a smaller field of view in Figure S3). The sources of these scaling factors are discussed in the next section.

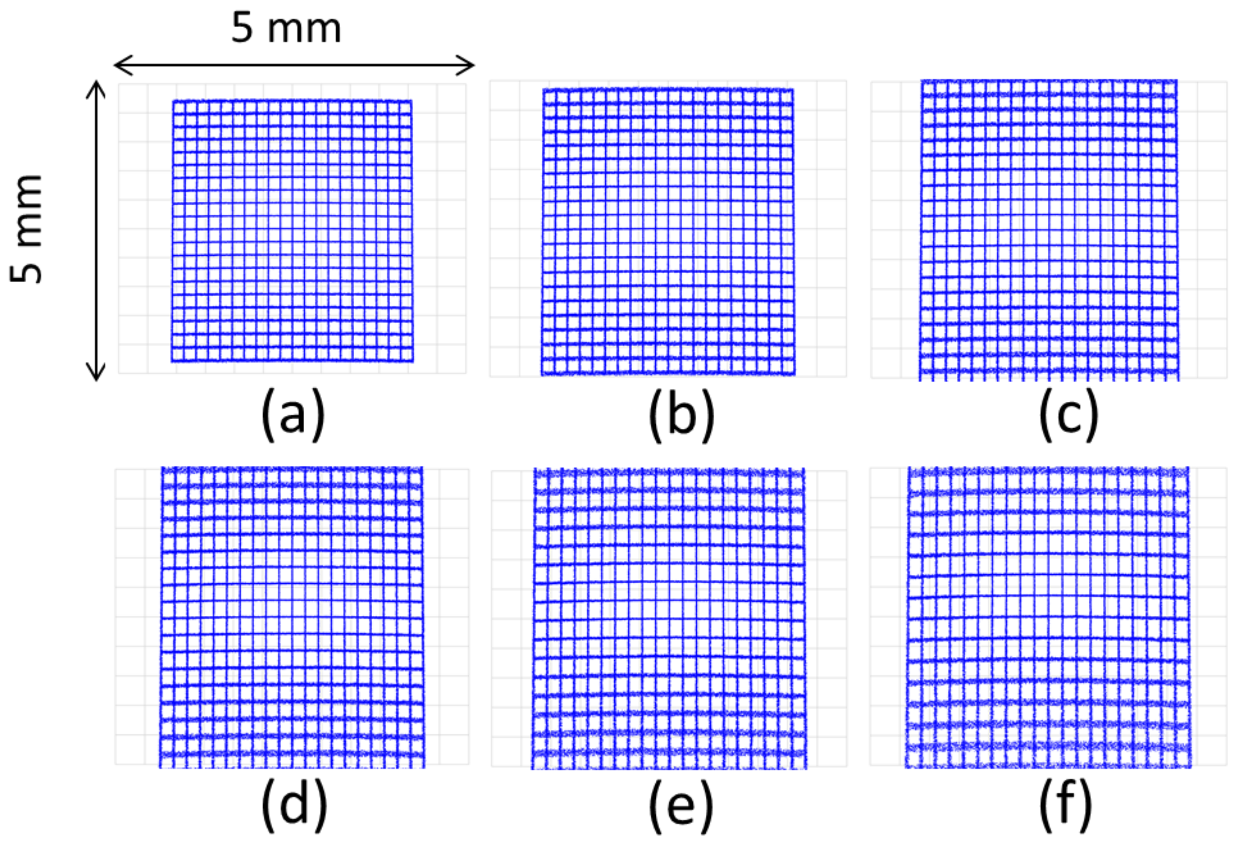


Figure S2: ZEMAX simulated images of a 4 mm × 4 mm grid with 200 µm divisions. Figures (a)-(f) correspond to the configurations 4-9 listed in Table 1.

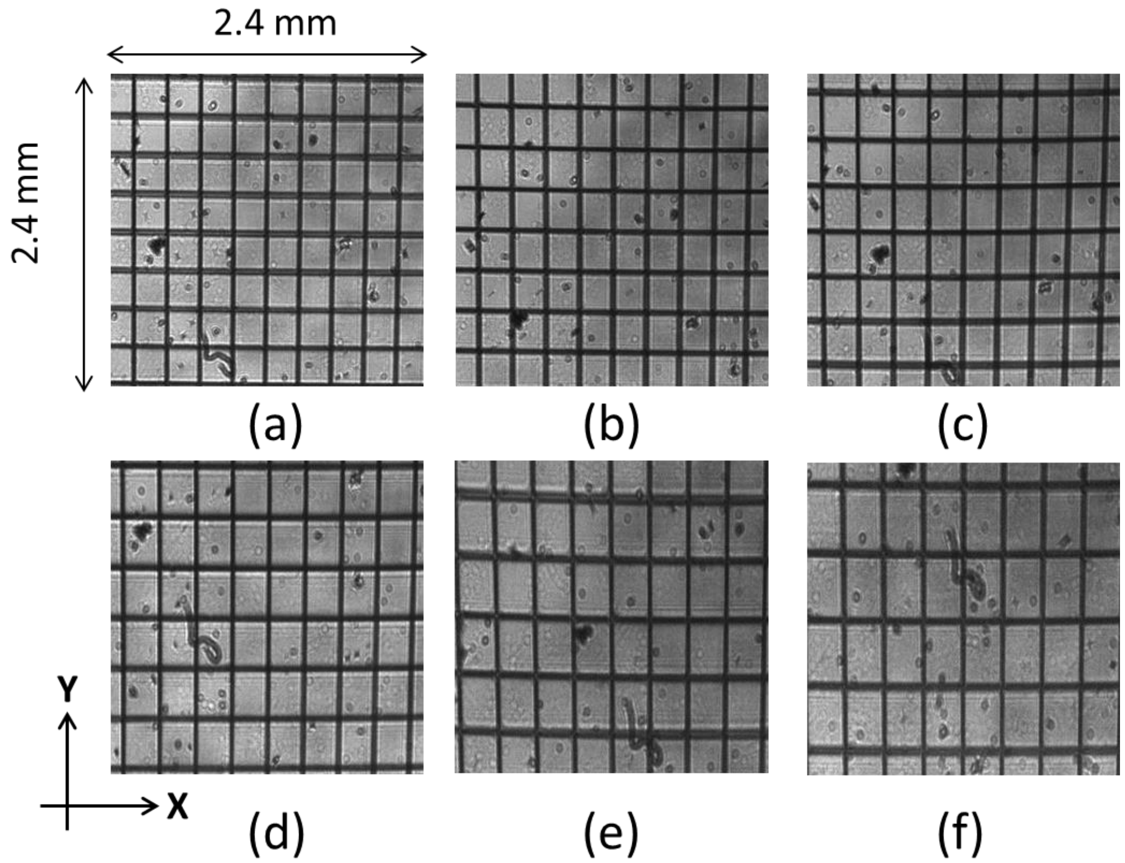


Figure S3: (a)-(f) Experimental images (400 × 400 pixels) of an opaque grid having a 200 µm pitch (Edmund Optics 58607) imaged from 50-100 mm off axis in 10 mm increments.

**Geometric projection**

As can be seen from Figure 2 in the main text, and Figure S1 and Table 1, the chief ray passing through the centre of the tube lens arrives at the detector plane at an incidence angle which increases with off-axis distance. As the image itself is formed normal to the chief ray (Figure S4), the detector plane therefore sees a geometric projection of the image. The image therefore appears stretched along the direction of off-axis displacement. In addition, the image moves through focus from one side of the detector plane to the other. The region over which the image is in focus is determined by the depth of focus of the parabolic mirror (approximately 5mm). Even at the highest incidence angle of 53°, the distance along the chief ray between the image formed at one side of the 2.4 mm detector and the other is given by 2.4 * sin(53°) = 1.44 mm and the entire image therefore remains inside the Rayleigh length of the parabolic focus.

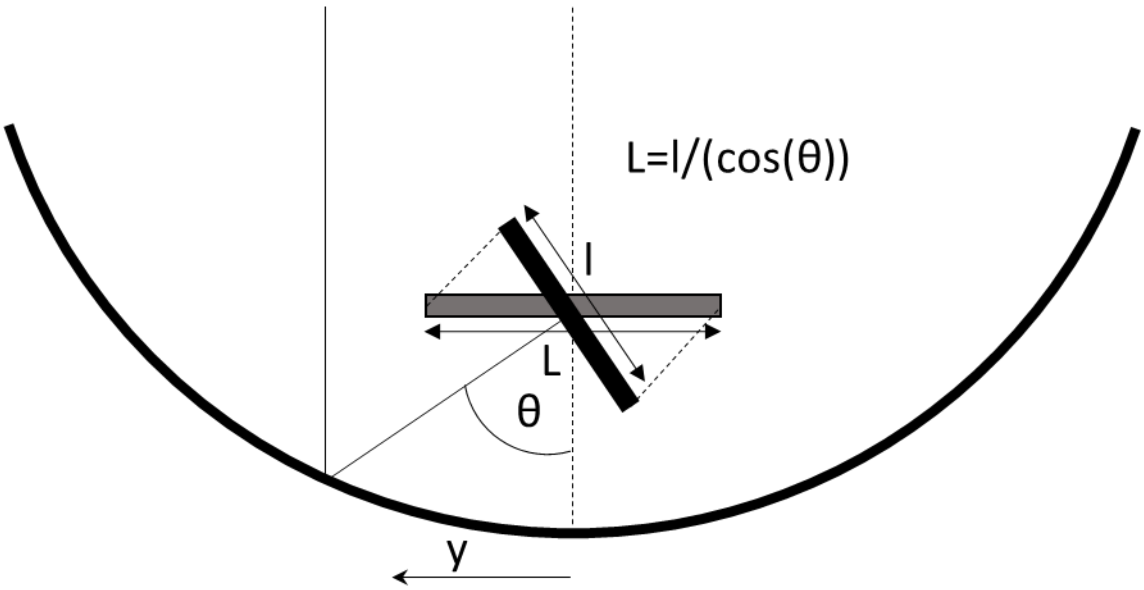


Figure S4: Model showing the effect of geometric projection.

The variation in the angle of incidence as a function of the off-axis distance was calculated using:

$$\theta = 2\,tan^{-1}\left(\frac{y}{2f}\right)$$

Where θ is the angle of incidence, *y* is the off-axis distance and *f* is the focal length of the parabolic mirror. This description of the elongation of the image was tested in simulations by rotating the detector plane by 33.4° for a 60 mm off-axis distance (Fig S5).

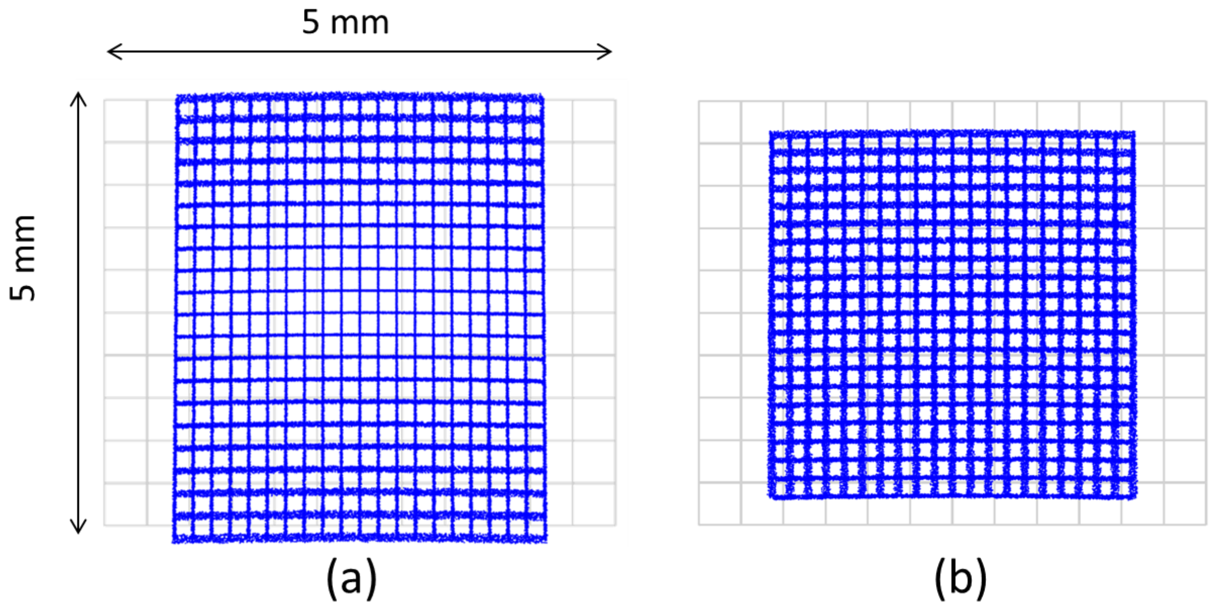


Figure S5: Simulation results illustrating the effect of geometric projection. (a) Image formed using an off-axis distance of 60 mm with the detector plane perpendicular to the optical axis of the parabolic mirror. (b) Image formed after rotation of the mirror by 33.4 ° in the direction of the light incident from the mirror. The geometric projection has been compensated.

**Global scaling**

In addition to geometric projection, the image also experiences scaling in both dimensions. As shown in , the distance between the mirror (the last focusing element) and the image increases with off-axis distance. The magnification is then given by the ratio of this distance (V) and the focal length of the tube lens.

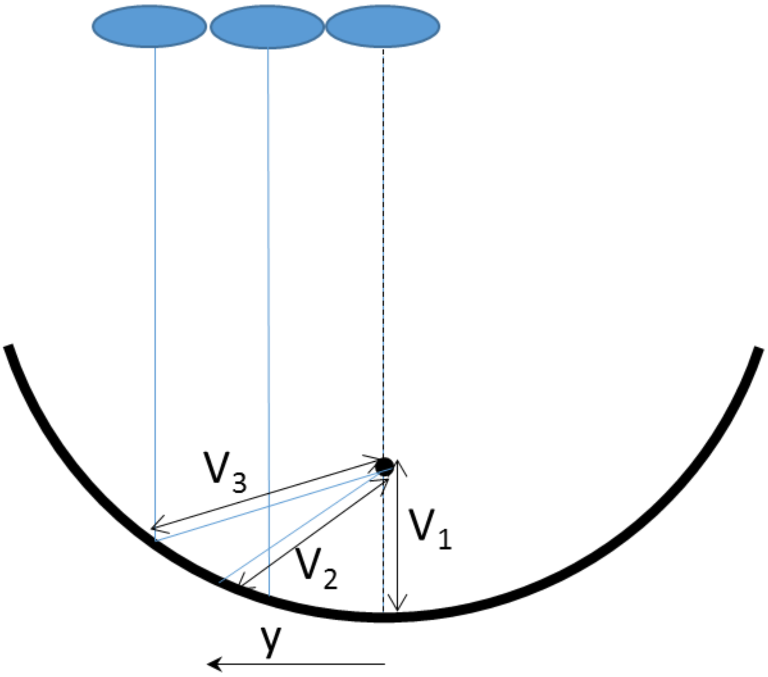


Figure S6: Sketch indicating the source of the global scaling. As the off-axis distance (y) increases, the distance between the mirror surface and the focal point also increases (V3>V2>V1). This leads to an increase in image magnification.

As the distance between the mirror surface and the focal point can be calculated precisely for a parabola, it allows us to calculate the magnification, *M*, as a function of off-axis distance, y, lens focal length, $f_L$ and mirror focal length $f_M$:

$$M = \frac{1}{f_L}\left\{y^2 + \left(f_M - \frac{y^2}{4f_M}\right)^2\right\}^{\frac{1}{2}}$$

Combining global scaling and geometric projection allows us to predict the scaling in both *x* and *y* dimensions and compare this with the scaling measured directly from the experimental images. This is shown in Figure S7. Note that errors arise in the measurement of the magnification in the experimental data as the magnification varies slightly across the image for the same reason as shown in Figure S6.

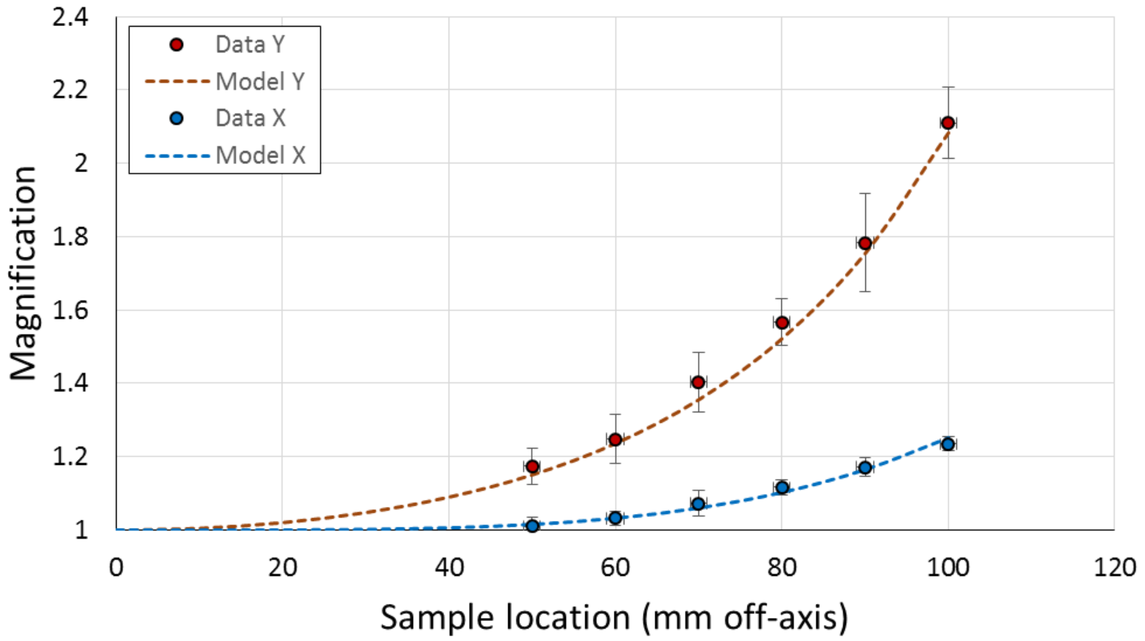


Figure S7: Plots showing the predicted (dashed lines) and actual (filled circles) magnifications for grid images at a variety of off-axis distances. Here the Y axis is taken to be parallel to the

**Image compensation**

As the sources of the image scaling are well known, they can also be compensated for any image without requiring a calibration step. So long as the off-axis distance and the focal lengths of the mirror and lens are known, the image magnification can be calculated and compensated. The image data shown in Figure S3 have been compensated using the above formulae and the results are shown in Figure S8. Note that in order to retain the full field of view, a large image (1280 × 1024 pixels) was first captured using a larger size detector. These images were then numerically compensated and the central 400 x 400 pixel region segmented. This allows a 1:1 comparison with the images acquired in Figure S3. In general, the field of view will be reduced as a consequence of the image scaling. The extent of this has been shown in Figure S9.

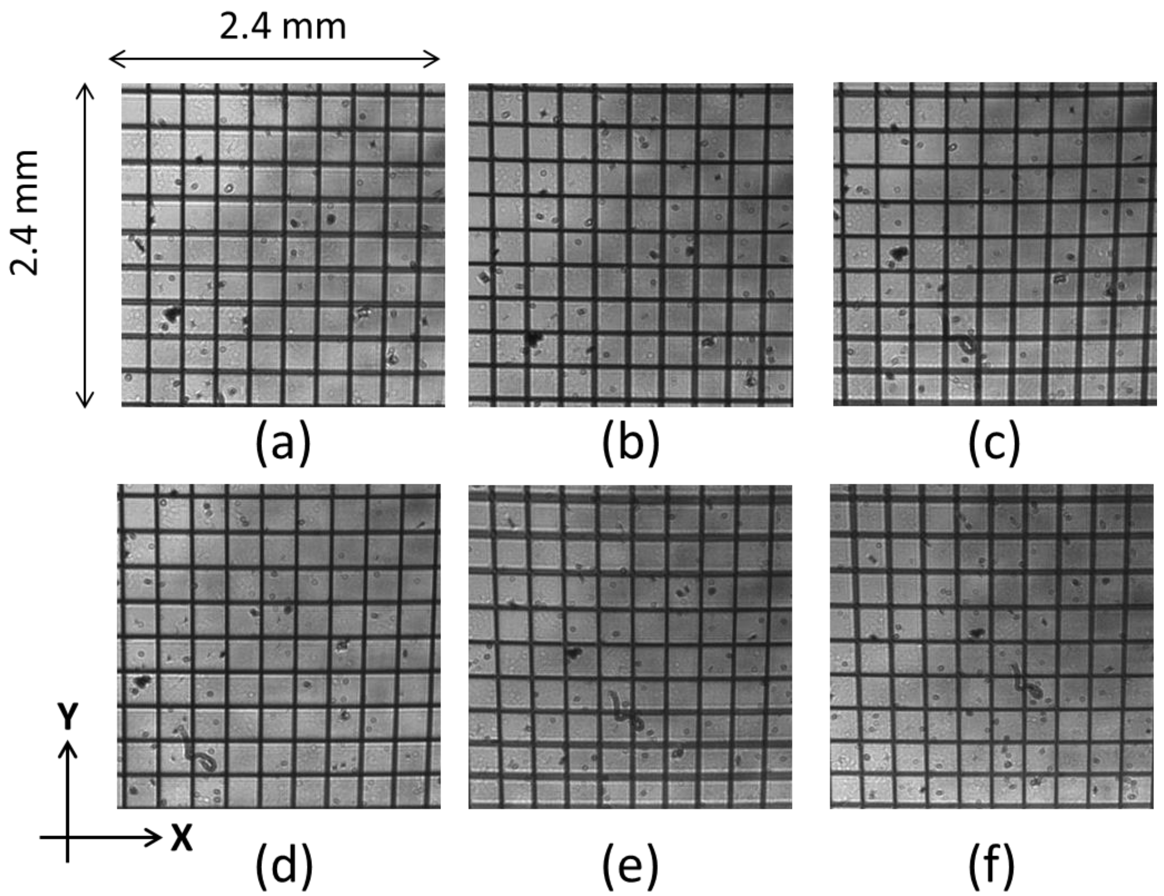


Figure S8: The same images as shown in Figure S3 after magnification compensation.

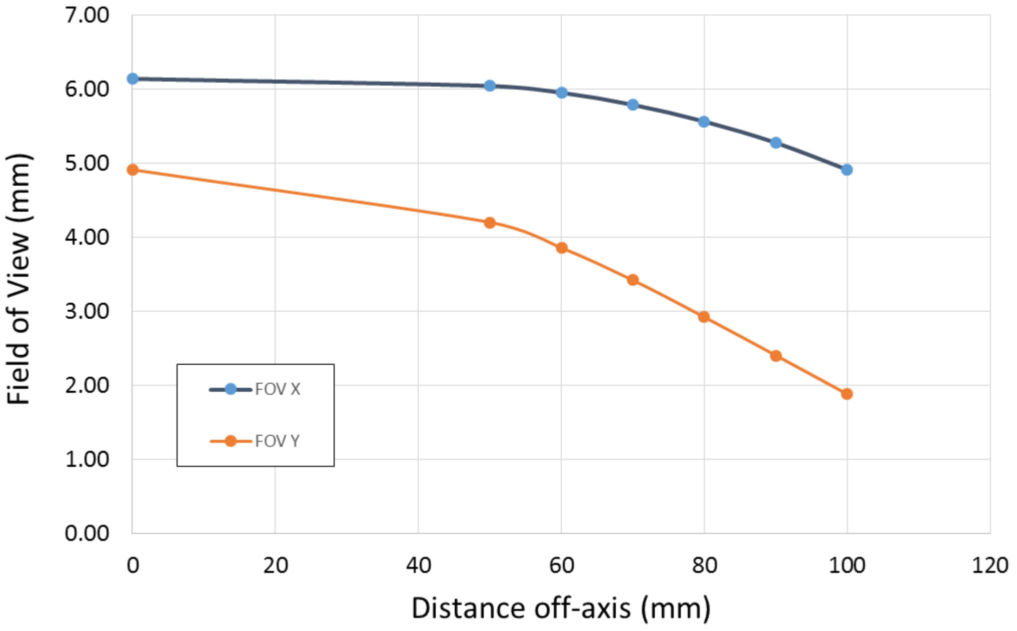


Figure S9: Graph indicating the reduction in the field of view as a function of off-axis distance for a 1280 × 1024 pixel detector.

A more quantitative comparison between Figure S3 and Figure S8 was performed in the Fourier domain. The Fourier Transform of each image was calculated and a line transect (in *x* and *y*) was taken through the first four orders either side of the zero order. This was performed for images acquired at 50-90 mm at 10 mm intervals. A comparison between the profiles before and after correction are shown for the *x* (Figure S10) and *y* (Figure S11) dimensions.

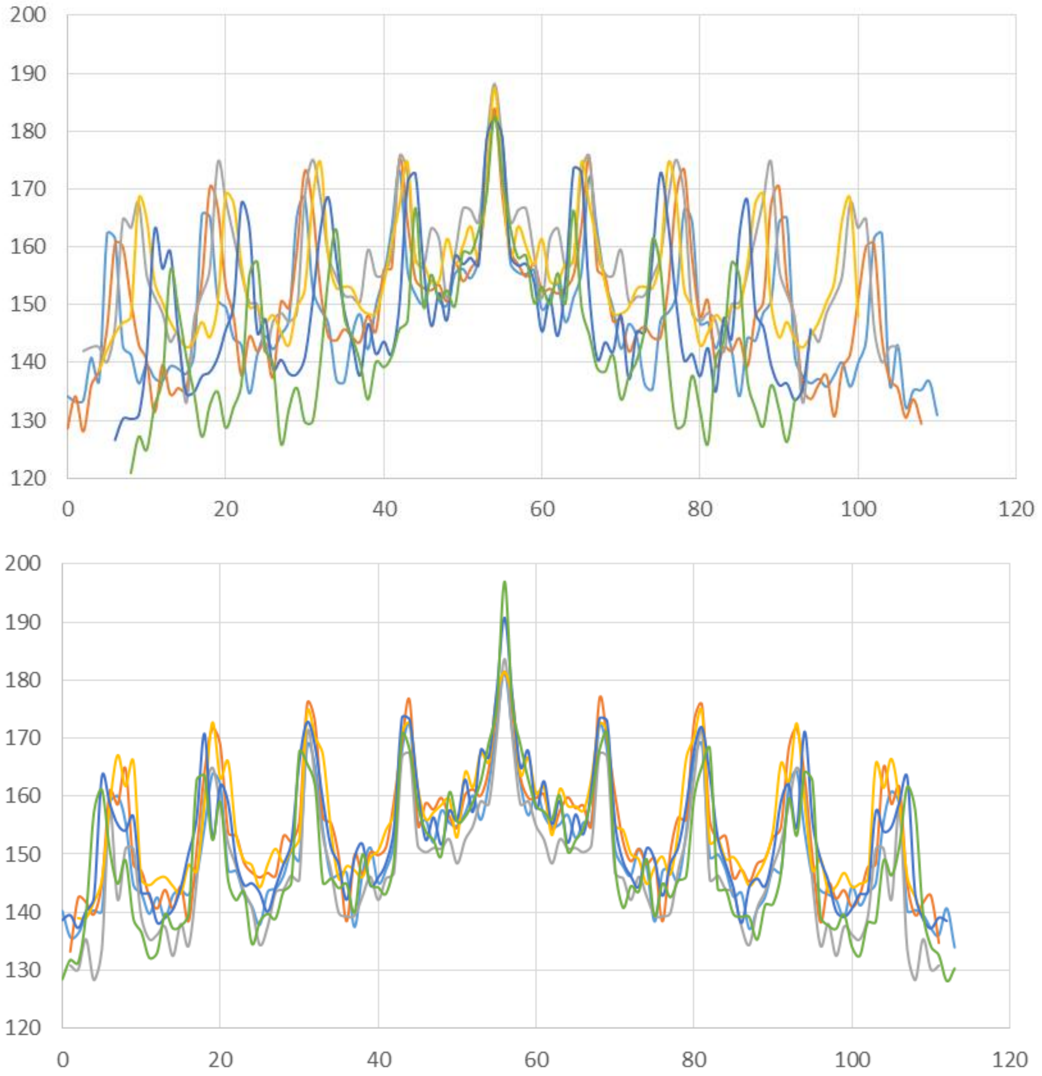


Figure S10: Line profiles through the *x*-dimension of the Fourier Transforms of experimental grid images. (Above) Line profiles for the images shown in Figure S3 and (below) those in Figure S8. Vertical axis: spectral power. Horizontal axis: Pixel value.

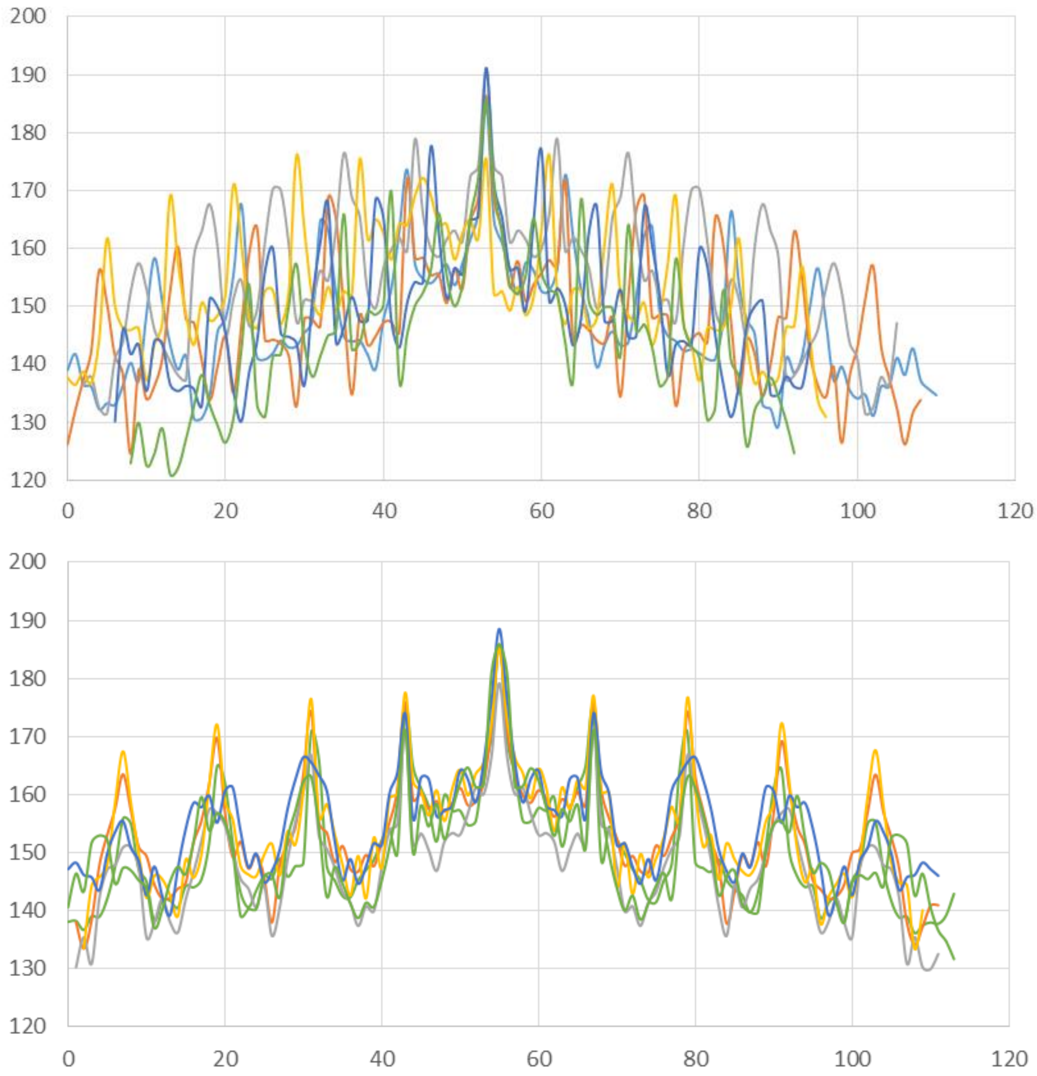


Figure S11: Line profiles through the y-dimension of the Fourier Transforms of experimental grid images. (Above) Line profiles for the images shown in Figure S3 and (below) those in Figure S8.

**Longer focal length mirrors**

One method to reduce the size of the magnification correction is to increase the focal length of the parabolic mirror. A longer focal length produces a shallower angle of incidence for the same axial offset. To maintain unitary magnification, the tube lens focal length would need to increase to match that of the mirror. This would then enable more samples to be imaged over a large range of offset values or the same number of samples to be imaged with a smaller magnification correction.
